# Cysteine-rich repeats trace past horizontal gene transfers in eukaryotes

**DOI:** 10.64898/2026.06.07.730663

**Authors:** Maria Daugavet, Victoria Dikaya, Natella Enukashvily, Stas Malavin, Maxim Rubin-Blum

## Abstract

Movement of genetic material between non-parental organisms, called horizontal gene transfer (HGT), is well recognized in prokaryotes but represents an underestimated force for the acquisition of novel traits in eukaryotes. The mechanisms of cross-domain HGT remain poorly understood despite numerous reports of its occurrence. Thus, progress in the field remains limited by the lack of targeted approaches for detecting HGT events. Using BLASTp search and sequence identity, we describe an HGT-derived protein from the chytridiomycete fungus *Neocallimastix californiae*. This protein has two cysteine-rich repeats (CysRReps) and other functional domains that are highly similar to those of prokaryotes. Based on the alignment of several CysRReps, we identified 859 additional eukaryotic proteins spanning 43 protein families, each with at least one match to bacterial, archaeal, or viral proteins. HGT-derived proteins with CysRReps belong to multiple eukaryotic taxa, some of which are well-known HGT models. Bacterial, archaeal, and viral proteins comprise the prokaryotic counterparts. Bacteria belong to 15 phyla, spanning a diverse array of physiologies, habitats, and lifestyles. Viral diversity is largely restricted to Caudoviricetes, suggesting a virus-mediated mechanism of DNA integration underlying these transfers. Although the function of CysRReps remains unclear, their association with HGT events across diverse eukaryotic and prokaryotic taxa suggests the existence of a universal molecular mechanism that facilitates gene transfer across phylogenetic boundaries.

## Introduction

Genetic novelty is a fundamental driver of evolutionary change. In addition to the mutational process and sexual reproduction, the movement of genetic material outside of parent-to-offspring inheritance, called horizontal gene transfer (HGT), represents an underestimated force in the acquisition of novel traits (Keeling and Palmer 2008). In prokaryotes, HGT can profoundly alter cellular physiology (Soucy et al. 2015), including the acquisition of genes conferring antibiotic resistance (Murray et al. 2022). HGT is not limited to exchange within one domain of life, as such events can cross domain boundaries (Boto 2014; Quispe-Huamanquispe et al. 2017; Cote-L’Heureux et al. 2022; Haimlich et al. 2024; Keeling 2024; Liu et al. 2024). In eukaryotes, HGT has likewise contributed to evolutionarily significant innovations, enabling, for example, the incorporation of novel amino acids into protein synthesis (Syvanen 2002), the acquisition of lysyl oxidase required for extracellular matrix cross-linking in multicellular organisms (Grau-Bové et al. 2015), the ability to build calcified skeletons (Jackson et al. 2011; Ettensohn 2014; Chess et al. 2025), or cellulose-based coverings (Sasakura et al. 2005; Inoue et al. 2019).

Several studies aimed to detect groups of eukaryotes more prone to DNA exchange with evolutionary-distant organisms. “Protists” seem to be good acceptors of foreign DNA because they lack a segregated germ line (Huang 2013). Among multicellular organisms, fungi are one of the groups for which multiple cases of HGT have been described (Fitzpatrick 2012; Ciach et al. 2024; Liu et al. 2024). Particular ecological niches or lifestyles may facilitate HGT, especially parasitic relationships and intimate symbiotic interactions between eukaryotic and prokaryotic partners (Nikoh et al. 2008, 2010; Klasson et al. 2009; Husnik and McCutcheon 2018; Ip et al. 2021), as increased physical proximity and prolonged cohabitation can enhance the likelihood of DNA exchange (Husnik and McCutcheon 2016; Bublitz et al. 2019). A comparison of multiple fungal genomes also showed that animal and plant parasites exhibit the highest estimated numbers of HGT-acquired genes among saprotrophs, symbionts, and parasites (Liu et al. 2024). In addition, desiccation was suggested as a factor in the susceptibility of bdelloid rotifers to exogenous DNA (Hespeels et al. 2014; Bininda-Emonds et al. 2016), but this was later debated in a systematic review of multiple taxa (Li et al. 2020). Intracellular bacteria or organelles of bacterial origin can serve as a readily available source of exogenous DNA (Stegemann et al. 2003). For example, when the mollusk *Elysia chlorotica* grazes on algae, it acquires functional plastids and incorporates photosynthesis-related genes into its own genome(Rumpho et al. 2008). At the end of this continuum, HGT from long-established endosymbiotic organelles, such as mitochondria and plastids, exemplifies a well-documented case of intimate genetic exchange (Timmis et al. 2004; Bock and Timmis 2008; Wang et al. 2022).

Detailed mechanisms of HGT have been studied in prokaryotic organisms and include conjugation (direct exchange of DNA between two cells), transformation (uptake of DNA from the environment), and transduction (virus-mediated DNA transfer) (Thomas and Nielsen 2005). Only a few cases of cross-domain HGT have been studied in detail, and the molecular mechanisms underlying DNA transfer and assimilation remain poorly described. One example involves plants and their root-knot bacterial symbionts, which use a type IV secretion system to deliver DNA across the eukaryotic plasma membrane (Gelvin 2003; Lacroix and Citovsky 2016). Once inside the plant nucleus, the transferred DNA can persist as an episome or integrate into the eukaryotic chromosome (Sheludko 2008; Pereman et al. 2019; Singer et al. 2022).

Despite extensive literature on cross-domain HGT, a limited understanding of the underlying mechanisms means most reported cases have been discovered incidentally, and systematic approaches for detecting such events remain limited (Irwin et al. 2022; Cote-L’Heureux et al. 2022). Targeted phylogeny-independent detection of HGT signatures is essential for uncovering the mechanisms of DNA transfer across domains of life. In this study, we investigate a putative HGT-derived protein from the chytridiomycete fungus *Neocallimastix californiae* and use a short amino acid hallmark within this protein as a search motif to identify multiple sequences across eukaryotic genomes. Using this approach, we detect cross-domain HGT events that have contributed functional protein sequences to diverse eukaryotic lineages. Although the mechanisms behind these transfers remain unclear, our findings provide new insight into the molecular signatures and evolutionary extent of cross-domain HGT.

## Methods

### Proteins of *N. californiae*

The proteins of *N. californiae* were taken from the genome assembly MCOG00000000 of *Neocallimastix californiae* strain G1 (Haitjema et al. 2017). We predicted functional domains and their protein families using InterProScan sequence search (https://www.ebi.ac.uk/interpro/search/sequence/), signal peptides by SignalP6.0 and subcellular localization with Phobious and InterPro (Kall et al. 2007; Teufel et al. 2022; Paysan-Lafosse et al. 2023). We searched for proteins similar to the *N. californiae* protein online using BLASTp against eukaryotes, bacteria, and viruses, using the NR database with an E-value threshold of 10^−4^. For graphical representation, we filtered the entries based on their score.

### Genetic context in bacteria and viruses

We identified prophage regions, positions of *attL* and *attR* recombination sites and protein-coding genes within the prophage region using Phaster (https://phastest.ca/) (Arndt et al. 2016). We obtained gene positions from bacterial and bacteriophage genomes in the GenBank entries JAMPGK000000000.1 and BK029377.1, respectively. An additional search was performed using standalone BLASTn (blast 2.15.0) for bacterial and viral contigs against a homemade database of well-described recombination sites extracted from GenBank records, filtered to be longer than 10 bp (Supplementary File 1).

### Cysteine-rich repeats containing proteins

We used the l-INS-i algorithm of MAFFT (v7.490) to align five CysRRep sequences from two *N. californiae* proteins (Supplementary File 2). We used this alignment to search the UniProtKB database with JackHMMER (v3.3.2) (Eddy 2011). For each hit protein, the taxonomic position of an organism was retrieved for further grouping. Each Eukaryotic protein sequence was then divided into regions according to CysRRep position, namely, the sequence before the first repeat starting from the N-terminus, sequence after the last repeat and sequence or sequences between repeats. We retained only regions longer than 100 amino acids for subsequent analysis. We performed searches for homologous sequences of those regions using MMSeqs2 (v.1d78ed30843f602cd73113e02639ce76bcb4b854) (Steinegger and Söding 2017) against the NR databases (release May 25 2025) of eukaryotes, bacteria, archaea, and viruses. We filtered the results using an E-value threshold of 10^−4^, and the best hit for each sequence region was recorded (Supplementary Files 3 and 4). We retained all eukaryotic sequence regions with hits to any prokaryotic protein and searched them against the Pfam database fasta files (v35.0) (Mistry et al. 2021) using MMSeqs2 (Supplementary File 5).

## Results and Discussion

### Cysteine-rich repeats may serve as recognizable markers of horizontally acquired proteins in *Neocallimastix californiae*

The previously described glycosyl hydrolase gene of *N. californiae* (GH, GenBank ID ORY85279.1) shows strong evidence of bacterial origin and contains recognizable cysteine-rich repeats (CysRReps), with conserved cysteine spacing Cx_6-9_Cx_5-7_Cx_8-11_Cx_7-8_CC (Daugavet et al. 2020). It has a predicted signal peptide at the N-terminus, targeting the protein for secretion. The mature protein begins with a pair of CysRReps followed by the glycosyl hydrolase domain. Apart from three related fungal sequences, the closest homologs this domain are bacterial proteins. Another CysRReps region follows again after the glycosyl hydrolase domain, although the second repeat is interrupted by an intron at the position where Cys-Cys amino acids could have been. The CysRRep region is followed by a peptidase domain whose closest homologue is likewise bacterial (Fig. 1A).

**Fig. 1.**
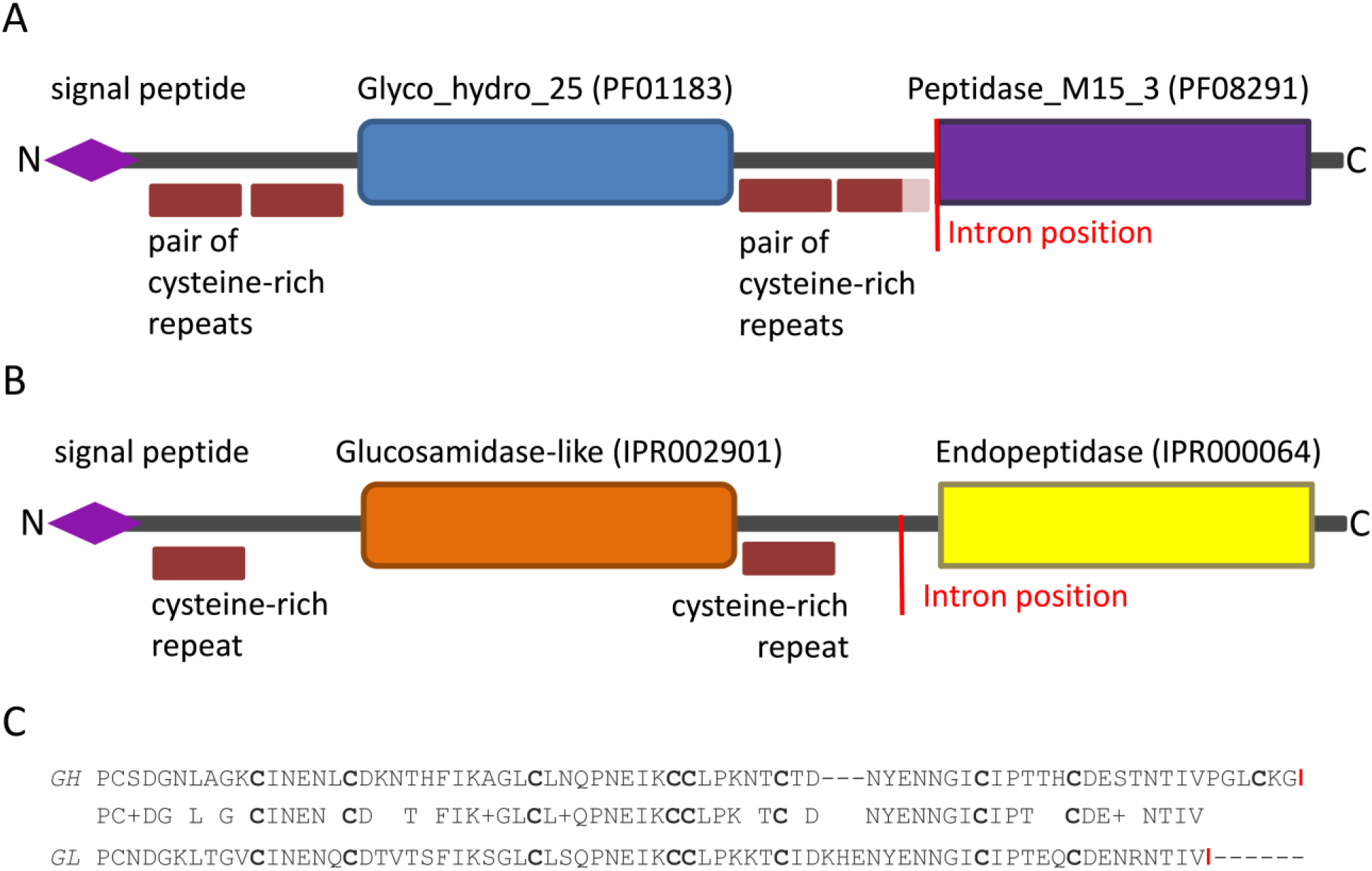
Domain architecture of two N. californiae proteins.**A** Glycoside hydrolase (GH) with two enzymatic domains interspaced by cysteine-rich repeats **B** glucosamidase-like (GL) protein with two enzymatic domains interspaced by cysteine-rich repeats **C** region of similarity in primary amino acid sequence between the two proteins. Conservative cysteines are bold, and red vertical lines indicate intron positions

The same structure but with a different primary sequence is characteristic of another protein from *N. californiae*, the glucosaminidase-domain-containing protein (GenBank ID ORY21964.1). It also has a predicted signal peptide at 1-25 aa at the N-terminus, targeting secretion (Fig. 1B; Supplementary Figure S1). We predicted all the remaining parts as non-cytoplasmic. Mature protein starts with one cysteine-rich repeat. Next is the enzymatic domain defined as glucosamidase-like (InterPro ID IPR002901). Hereafter, we will refer to this protein as GL (glucosamidase-like). After the glucosaminidase-like domain, CysRRep appears again, followed by an endopeptidase domain (InterPro ID IPR000064) (Fig. 1B). The gene contains a single intron located upstream of the coding region of the endopeptidase domain (Fig. 1B; Supplementary Figure S1).

In both proteins, CysRReps precede enzymatic domains. Nevertheless, the only region showing primary amino acid sequence similarity between the two proteins is CysRRep, with an adjusting sequence located before the second enzymatic domain and extending up to the intron position (Fig. 1C, red lines). For the previously described gene encoding a glycoside hydrolase, we proposed a chimeric combination of eukaryotic and laterally transferred prokaryotic parts (Daugavet et al. 2020). In this paper we aimed to identify the ancestry of the GL protein in *N. californiae* to determine whether the composition of protein parts could predict the gene’s origin.

### GL protein domains show stronger similarity to prokaryotic than to eukaryotic sequences

Assuming that a eukaryotic gene should have a eukaryotic predecessor, we searched the *N. californiae* GL protein against the NR database of eukaryotic sequences (excluding Neocallimastix) using BLASTp. We found very few proteins with similar regions (Fig. 2A). A short sequence from *Vanrija albida* (Basidiomycota) overlaps with the first CysRRep. In contrast, proteins from other eukaryotes overlap only with the glucosaminidase-like or endopeptidase domains and show no similarity to the CysRReps. These eukaryotic homologs occur only in crustaceans, insects, plants, and several protist groups (Supplementary Table T1 and File T1). Organisms possessing similar proteins with glucosaminidase-like or endopeptidase domains are unlikely to represent the direct ancestor of the GL gene, as they belong to distantly related phylogenetic groups. This patchy distribution pattern is characteristic for horizontally acquired genes (Fitzpatrick et al. 2008; Aubin et al. 2021).

**Fig. 2.**
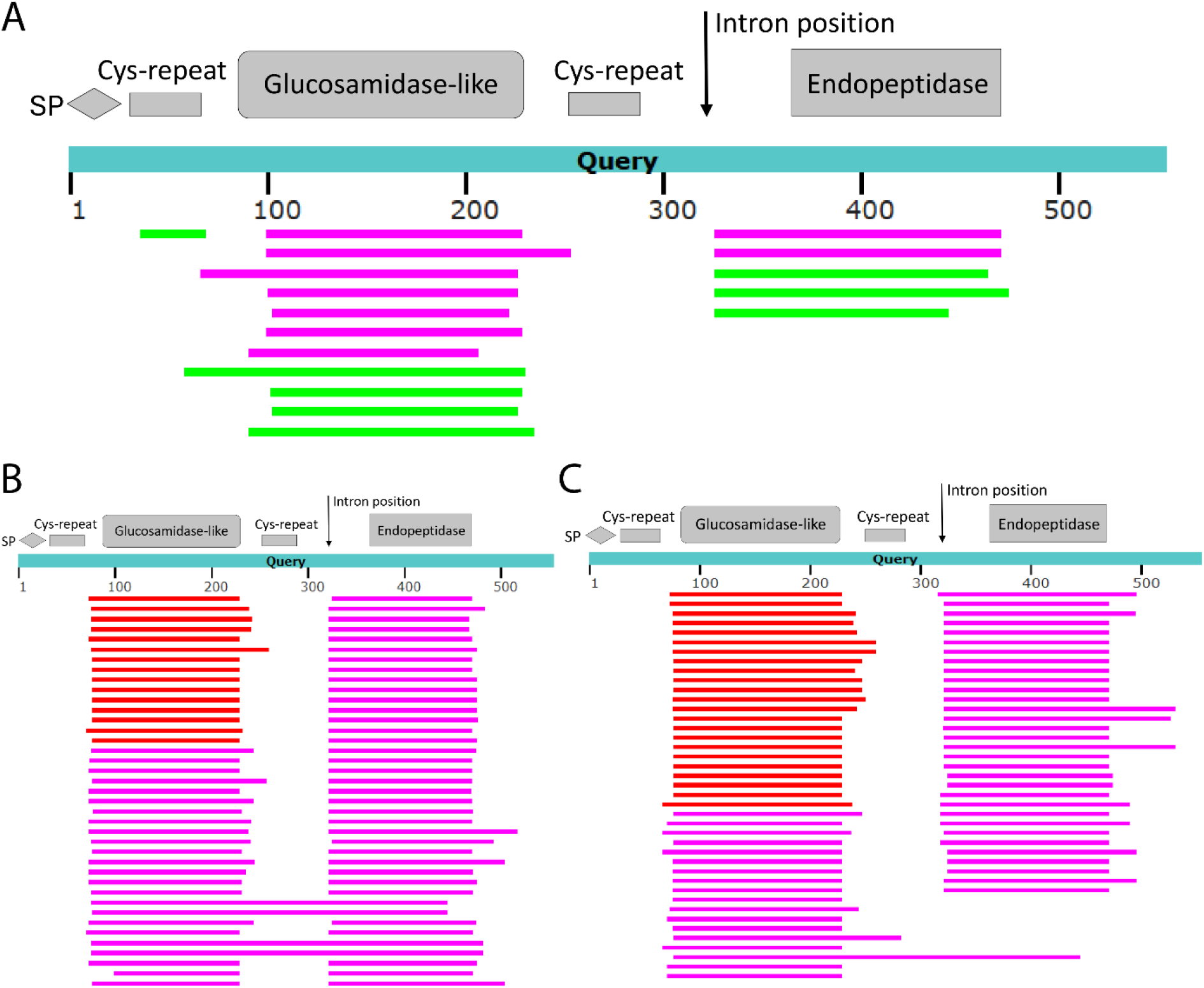
Representation of the BLAST search for the glucosamidase-like protein of *N. californiae*. Hits with **A** eukaryotic proteins **B** bacterial proteins and **C** viral proteins. Colors of the bars indicate alignment score (50-80, green; 80-200, purple; >200, red)

BLASTp search against bacterial sequences returned more than 1000 hits with 28–67% identity, yet none covered the full query sequence. Most bacterial matches correspond only to one of the two enzymatic domains and lack the CysRReps. For the glucosaminidase-like domain, similarity to bacterial sequences exceeds that to non-fungal eukaryotic sequences (Fig. 2B; Supplementary Table T2). When we searched for similar viral sequences using BLASTp, we obtained 735 hits with sequence identities ranging from 22% to 67%. Most of the hits overlapped only with one of the predicted domains (Fig. 2C; Supplementary Table T3). We therefore infer that the glucosaminidase-like and endopeptidase domains are more closely related to prokaryotic (bacterial or viral) sequences than to any eukaryotic homologs.

We next sought to identify the closest relative of a prokaryotic organism that contributed the sequence later integrated into the genome of *N. californiae*. We queried the NR database with BLASTp using amino acid region Q76–D228 of *N. californiae*, corresponding only to the predicted glucosaminidase-like domain (Supplementary Figure S1). Excluding hits with 100% identity to the full-length *N. californiae* sequence, the best match was a glucosaminidase domain-containing protein from a Lachnospiraceae bacterium (isolate NALB_7863_7; GenBank ID: MCM1218142.1), with 67% identity and an E-value of 10^−67^ (Fig. 3A). This isolate originates from the gut microbiome of the wild rodent *Neotoma albigula*. The second-best hit was a flagellum-specific muramidase from an unclassified Caudoviricetes bacteriophage of the human gut microbiome (GenBank ID: DAV93767.1), with 66% identity and an E-value of 5 × 10^−68^ (Fig. 3B).

**Fig. 3.**
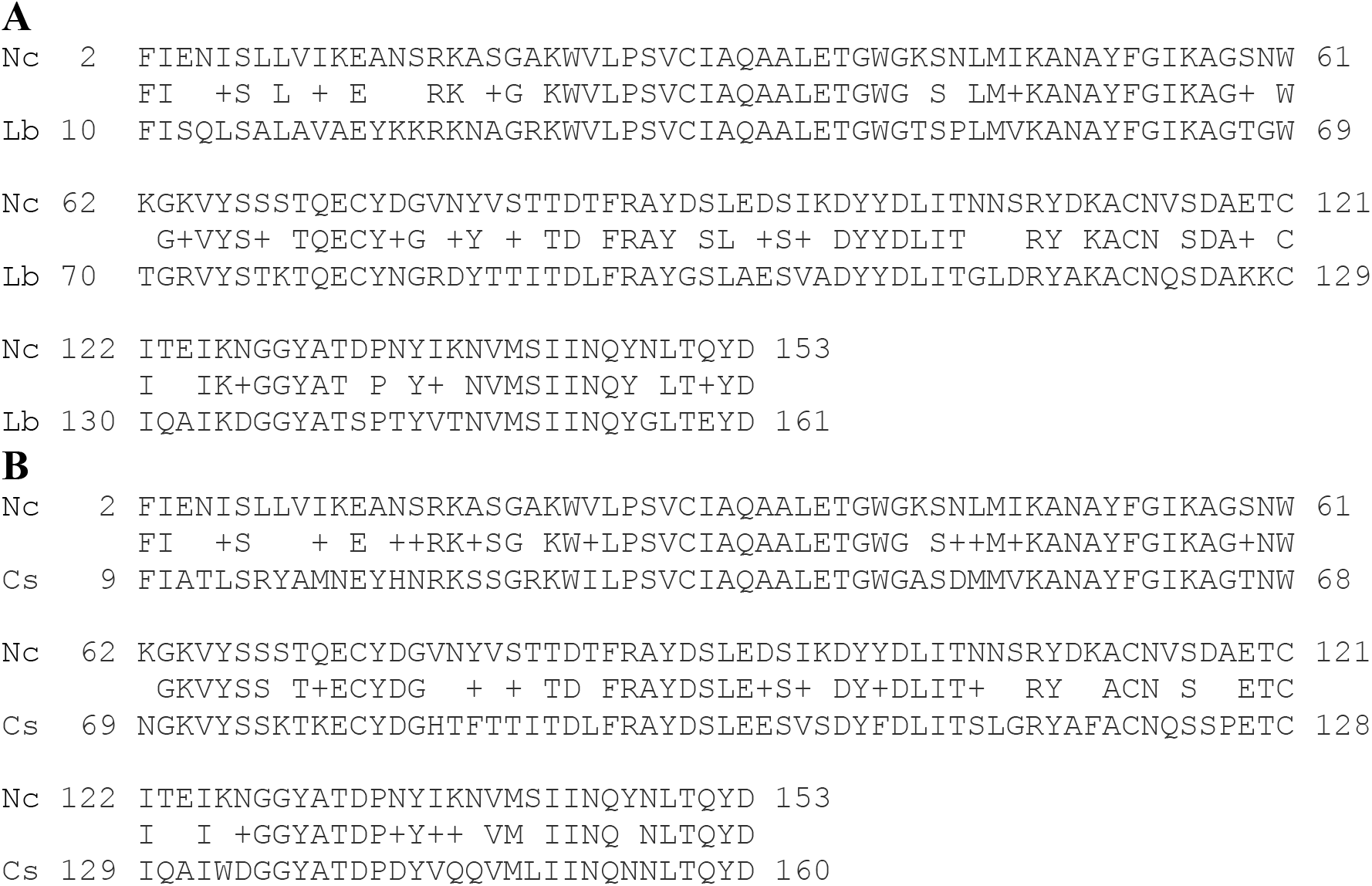
Alignments of *N. californiae* sequence corresponding to predicted glucosamidase-like domain. Alignment with **A** glucosaminidase domain-containing protein of Lachnospiraceae bacterium protein (MCM1218142.1) and **B** flagellum-specific muramidase of Caudoviricetes sp. virus (<u>DAV93767.1</u>). Nc - *N. californiae*, Lb - Lachnospiraceae bacterium, Cs - Caudoviricetes sp.

To evaluate the possibility of independent integration events, we conducted a separate search for homologs of the GL protein’s endopeptidase domain. We used the isolated amino acid region V367-Y463 (Supplementary Figure S1) as a BLASTp query. Excluding *N. californiae* hits, the best match was a Lachnospiraceae bacterium sequence (isolate C.4_2; GenBank ID: MCD8119760.1), with 57% identity and an E-value of 4 × 10^−30^ (Supplementary Figure S2). This isolate originates from the gut microbiome of a juvenile koala (*Phascolarctos cinereus*). We also found several significant matches among viral proteins, with the best hit corresponding to a CW7 repeat protein (DAT38862.1) of a Caudovoricetes virus from the human gut microbiome, showing 54% identity and an E-value of 2 × 10^−25^ (Supplementary Figure S3). Notably, although the *N. californiae* strain G1 was isolated from the goat gut, the closest homologous sequences were found in bacteria from the gut microbiomes of other animal hosts. Likewise, the identified bacteriophages are associated with the human gut microbiome. Therefore, the detected bacterial and viral sequences likely represent relatives of the original donor organisms rather than the direct sources of the DNA that gave rise to the *N. californiae* GL protein.

### No evidence supports a prokaryotic origin for CysRRep

Since the two enzymatic domains of the GL protein have close homologs in Lachnospiraceae, we hypothesized that the CysRReps might likewise be of bacterial origin. To find out, we aligned 5 CysRReps: three from the *N. californiae* glycoside hydrolase protein described previously and two from the *N. californiae* GL protein described here (Supplementary File 2). The JackHMMER search against UniProtKB with no taxonomic restrictions identified multiple eukaryotic hits, 35 bacterial hits (27 Myxococcota, 1 Pseudomonadota, 1 Actinomycetota, 1 Candidatus Levyibacteriota, and 5 species with no phylum defined) and no matches with virus proteins (Supplementary Table T4). None of the bacterial hits matched Bacillota, to which the *Lachnospiraceae* bacteria belong. Hence, we cannot consider those repeats as predecessors of the *N. californiae* cysteine-rich coding sequence.

The longest bacterial hit of GL protein overlaps with both glucosaminidase-like and endopeptidase domains with an overall identity of 33% and an E-value of 6 × 10^−41^. It contains a highly gapped region corresponding to the middle CysRRep, extending approximately to the position of the intron (Supplementary Figure S4). That suggests the insertion of CysRReps after the transfer of a bacterial DNA fragment into the eukaryotic genome.

### Prophage context of homolog genes indicates phage-mediated transfer

To investigate the mechanism of foreign DNA integration for the GL protein, we analyzed the genetic context surrounding the transferred regions in the closest prokaryotic organisms. The bacterial homolog of the glucosaminidase-like domain is located on contig JAMPGK010000058.1, with a site-specific integrase 640 bp upstream and a phage holin family protein downstream (Fig. 4A). Both proteins are characteristic of bacteriophage genomes. The entire contig represents an incomplete prophage region that includes the flanking recombination sites AttR and AttL. This finding suggests that the bacterial sequence similar to the eukaryotic one does not inherently belong to the bacterial genome but instead reflects phage integration into the bacterial chromosome. A homolog of the glucosaminidase-like domain in the bacteriophage genome Caudoviricetes sp. (GenBank: BK029377.1) corresponds to a muramidase gene. In the phage genome, this gene overlaps two hypothetical protein-encoding genes of unknown function and lies 406 bp downstream of a site-specific integrase gene (Fig. 4B).

**Fig. 4.**
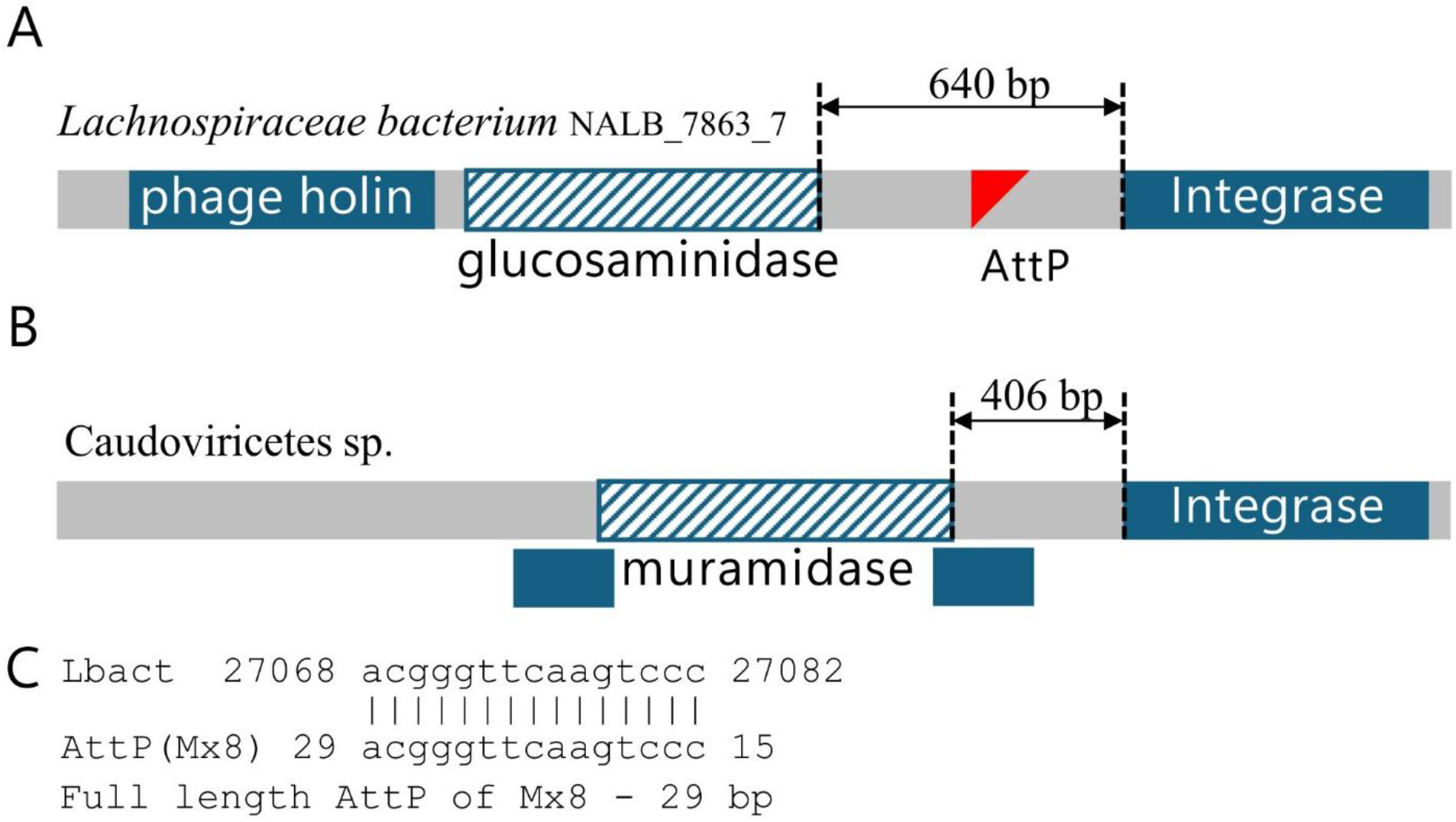
Genetic context of glucosaminidase-domain-containing protein homologs of N. californiae. **A** glucosamidase gene of Lachnospiraceae bacterium isolate NALB_7863_7 and **B** muramidase gene of Caudoviricetes sp. bacteriophage (GenBank ID: DAV93767.1), **C** alignment of Lachnospiraceae bacterium contig sequence (Lbact) with Myxococcus phage Mx8 recombination site (AttP(Mx8))

Canonical bacteriophage integration into bacterial chromosomes occurs via site-specific recombination, making the recombination site a useful marker of the integration breakpoint. In phages, AttP recombination sites often occur next to the integrase gene (Thorpe and Smith 1998; Julien 2003). Hence, we examined the regions between the integrase genes and the glucosaminidase-like domain homologs in the bacteriophage Caudoviricetes sp. genome and in the prophage region of a Lachnospiraceae bacterium for previously annotated AttP sequences. In the bacterial chromosome, we detected similarity to half of the AttP site of the *Myxococcus* phage Mx8 (GenBank ID D86464.1) (Fig. 4C). In the Caudoviricetes sp. genome, we found no AttP sequence in the region separating the muramidase and integrase genes.

The closest bacterial homolog of the endopeptidase domain occurs on contig JAJQBQ010000044.1 of the Lachnospiraceae bacterium isolate C.4_2 and lies within an intact prophage region. No integrase gene occurs within this prophage, suggesting potential loss during prophage degradation. The recombination sites *attL* and *attR* flank the prophage region but map to a distant genomic location relative to the predicted endopeptidase-coding gene (Supplementary File 6). In the Caudoviricetes phage genome, the chromosome (BK046062.1) harboring an endopeptidase domain homolog lacked an integrase nearby. In both closest sequences from bacterial chromosomes, the homologous genes lie within the prophage regions. This finding suggests that the Lachnospiraceae bacterium is unlikely to be the direct donor of foreign DNA to the *N. californiae*. We also note that the ancestral glucosaminidase-like gene lies close to a putative recombination site, supporting a bacteriophage-mediated integration event.

### Cysteine-rich repeats link eukaryotic genes to prokaryotic sequences

At least two *N. californiae* proteins containing CysRReps harbor genomic regions of putative prokaryotic origin. To assess whether this association extends beyond *N. californiae*, we analyzed a broader dataset of eukaryotic proteins containing CysRReps. By search in UniProtKB database, with the CysRReps alignment, we found 1246 protein matches with E-values < 1 × 10^−4^, belonging to 339 species (324 eukaryotes, 13 bacteria, 2 archaea; Supplementary File 7). We extracted sequences longer than 100 aa from these proteins, corresponding to the parts of the proteins that exclude CysRReps: 96 N-terminal sequences, from the start to the first repeat, 849 C-terminal sequences, after the last repeat to the end of the protein, and 18 sequences between repeats (Supplementary Files 8, 9, 10, respectively). This uneven distribution indicates that CysRReps occur far more often at the N-terminus than at the C-terminus, while long (>100 aa) intervening regions between repeats remain rare. The MMSeqs2 searches of each sequence resulted in 859 proteins having at least one hit to a non-eukaryotic protein (Fig 5A, Supplementary Table T5). More than 80% of the sequences located on the C-terminal side relative to CysRRep showed significant similarity to bacterial proteins.

**Fig. 5.**
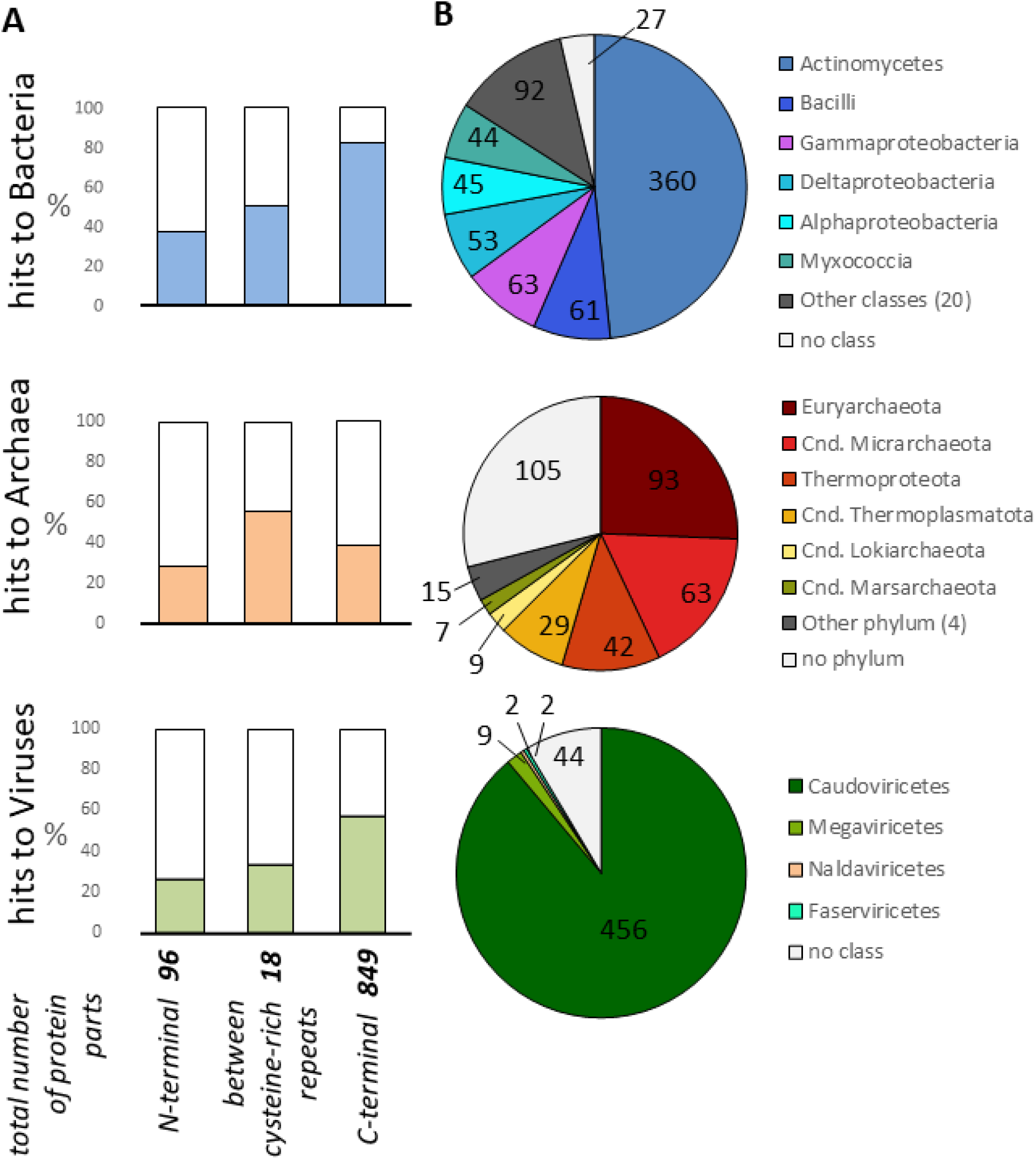
Protein parts with reliable similarity to different non-eukaryotic proteins. **A** Histograms show the fraction of all sequences that exhibit reliable similarity to proteins from different non-eukaryotic groups (blue, bacteria; orange, archaea; green, viruses; blank, no similarity). **B** Pie charts show the taxonomic distribution of species that possess those proteins

Our results suggest that eukaryotic proteins with CysRReps are similar and presumably share ancestry with those from taxonomically distinct bacteria and archaea. We asked which prokaryotic groups most frequently participate in CysRReps-associated HGT. Actinomycetes accounted for nearly half of all bacterial hits among the queries we found above, with others including Bacilli, Gammaproteobacteria, Deltaproteobacteria, Alphaproteobacteria, and Myxococcia (Fig 5B). In total, 26 bacterial classes were present. Among Archaea, Euryarchaeota, Candidatus Micrarchaeota, and Thermoproteota accounted for nearly half of the diversity. Viral diversity was largely restricted to Caudoviricetes, accounting for 89% of all hits, followed by Megaviricetes, Naldaviricetes, and Faserviricetes, as well as 23 unclassified phages. Taken together, the broad taxonomic distribution of homologs and the strong dominance of Caudoviricetes suggest that for CysRReps-containing proteins bacteriophage-mediated transfer is a major pathway linking prokaryotic gene pools with eukaryotic genomes.

### Widespread occurrence of cysteine-rich repeat proteins across eukaryotic lineages

We next examined the taxonomic distribution of these proteins to determine whether they occur in a single species, within a specific genus, or across broader taxonomic groups. We identified numerous eukaryotic proteins containing CysRReps that show signatures consistent with DNA exchange with bacteria and viruses. Proteins with CysRReps occur in four fungal phyla, nine metazoan phyla, one plant, and one algal phylum. Five proteins belong to *Spumella elongata*, a unicellular stramenopile. Within Fungi, Ascomycota contain the highest number of these proteins, and Rotifera dominate among Metazoa (Fig. 6A). We also assessed taxonomic diversity based on the number of species that contain CysRRep proteins rather than on protein counts. This analysis shows that these proteins occur across a wide range of eukaryotic lineages (Fig. 6B).

**Fig. 6.**
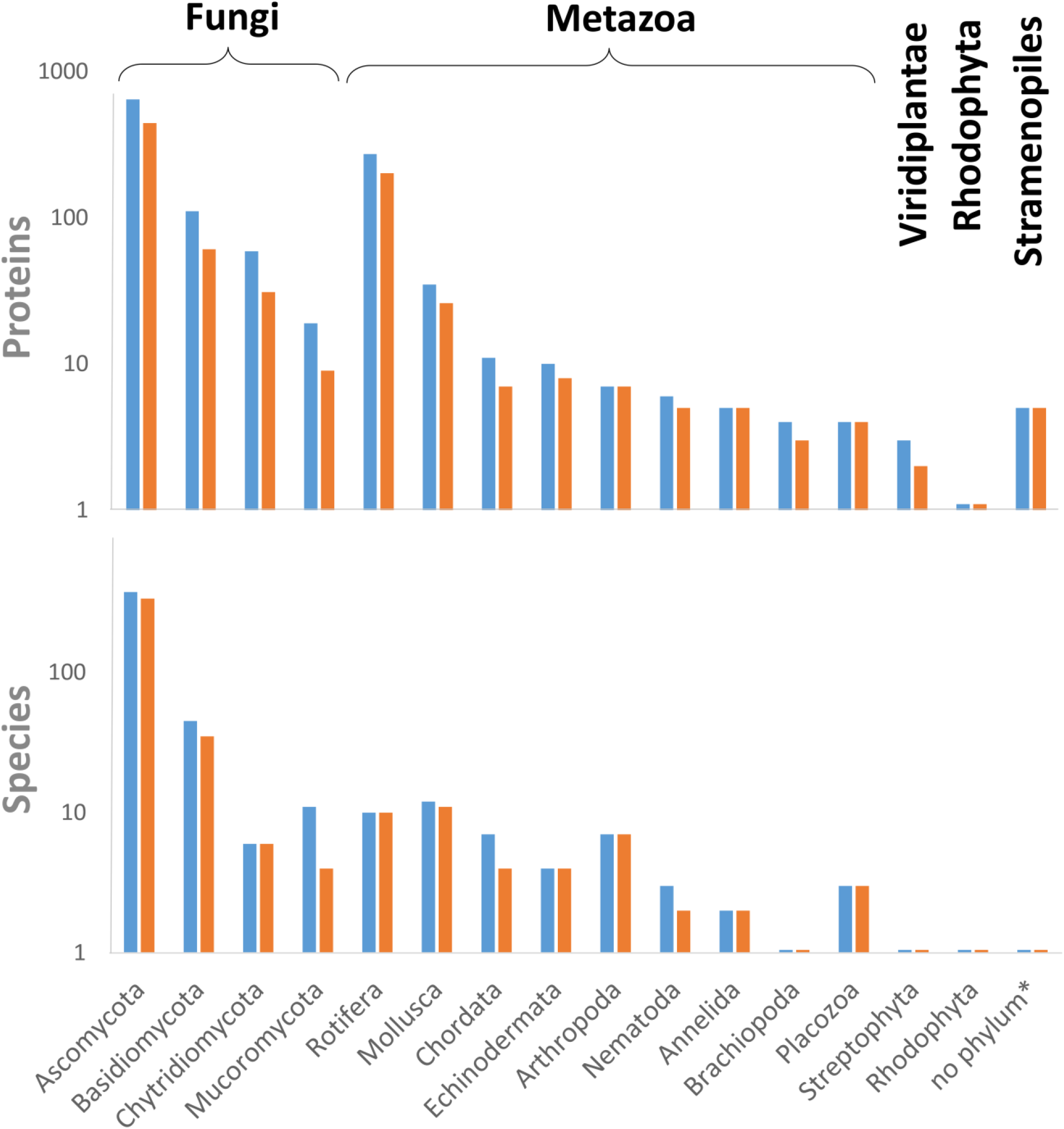
Distribution of proteins with cysteine-rich repeats (CysRReps). A Number of proteins with CysRReps in different groups of eukaryotes, total number - blue bars, only proteins with significant similarity to non-eukaryotic protein – orange bars. B Number of different species in different groups of eukaryotes, possessing proteins with CysRReps. Blue bars show the total number of species, and orange bars show only in cases where species have proteins with significant similarity to non-eukaryotic proteins

Across both analyses, the number of proteins with non-eukaryotic hits closely follows the total number of proteins containing CysRReps. The counts are equal or slightly lower, demonstrating that some proteins carry the CysRRep but show no detectable similarity to bacterial, archaeal, or viral sequences. However, a substantial fraction of these proteins display significant similarity to non-eukaryotic proteins, which we interpret as evidence of shared ancestry. This relationship remains consistent across the different eukaryotic groups examined.

### Diverse protein families support multiple independent HGT events

To test whether protein parts found across diverse eukaryotic clades originate from a single ancestral protein group, we determined their affiliation with protein families. We selected only those sequences that show similarity to non-eukaryotic proteins and compared them with the Pfam database of protein families. In total, 43 protein families were detected (Table 1). There is no obvious connection to inherently mobile elements like transposases (Schaack et al. 2010)—we found only one protein with a Tc1 transposase catalytic domain (PF13358) and a DNA recognition domain (PF01498). Frequencies of protein families were very similar, with no relation to proteins with hits to Bacteria, Archaea, or Viruses. We consider that the maximum number of different protein families we can observe for sequences on the C-terminal side is due to their greater abundance. The only exception to this rule is chitinase class I, which is more common at the N-terminal side. The total number of protein families covered in this analysis indicates that several ancestral sequences from different families independently participated in HGT.

**Table 1.** Protein families of sequence parts resembling bacterial, archaeal, or viral proteins. Counts are shown for sequences located N-terminal to the CysRReps (N), between repeats (M), and C-terminal to the CysRReps (C). Color indicates frequency: yellow (1-10), orange (11-40), and red (>40)

| Protein families in Eukaryotic sequence | Bacterial hits |  |  | Archaeal hits |  |  | Viral hits |  |  |
| --- | --- | --- | --- | --- | --- | --- | --- | --- | --- |
|  | N | M | C | N | M | C | N | M | C |
| Phage_lysozyme |  |  | 78 |  | 1 | 78 |  | 1 | 79 |
| D-alanyl-D-alanine_carboxypeptidase |  |  | 60 |  |  | 61 |  |  | 39 |
| N-acetylmuramoyl-L-alanine_amidase |  |  | 37 |  | 6 | 35 |  |  | 34 |
| NlpC/P60_family |  |  | 31 |  |  | 17 |  |  | 26 |
| Glycosyl_hydrolases_family_25 |  | 1 | 15 |  | 1 | 14 |  | 1 | 3 |
| Protein_of_unknown_function_(DUF3238) |  |  | 11 |  |  | 7 |  |  |  |
| Stanniocalcin_family | 2 |  | 6 |  |  | 3 |  |  |  |
| Amiloride-sensitive_sodium_channel |  |  | 5 |  |  |  |  |  | 1 |
| Destabilase | 2 |  | 5 |  |  | 4 |  |  | 5 |
| Aminopeptidase_P,_N-terminal_domain |  |  | 3 |  |  | 3 |  |  | 3 |
| Glycosyl_hydrolase-like_10 |  | 1 | 3 |  |  | 2 |  |  | 2 |
| Metallopeptidase_family_M24 |  |  | 3 |  |  | 3 |  |  | 3 |
| Nematode_trypsin-6-like_family |  |  | 3 |  |  | 3 |  |  | 3 |
| Peptidase_family_M23 |  |  | 3 |  |  | 3 |  |  | 3 |
| Phage_tail_lysozyme |  |  | 3 |  |  |  |  |  | 3 |
| Trypsin |  |  | 3 |  |  | 3 |  |  | 3 |
| Cell_Wall_Hydrolase |  |  | 2 |  |  | 1 |  |  | 1 |
| C-type_lysozyme/alpha-lactalbumin_family |  |  | 2 |  |  | 1 |  |  | 1 |
| Domain_of_unknown_function_(DUF1986) |  |  | 2 |  |  | 2 |  |  | 2 |
| Glycosyl_hydrolase_108 |  | 1 | 2 |  |  | 1 |  |  | 1 |
| Peptidase_M15_ |  | 1 | 2 |  |  | 2 |  |  | 2 |
| Prokaryotic_phospholipase_A2 |  |  | 2 |  |  |  |  |  |  |
| Alpha-L-rhamnosidase_N-terminal_domain |  |  | 1 |  |  | 1 |  |  | 1 |
| Astacin_(Peptidase_family_M12A) |  |  | 1 |  |  | 1 |  |  |  |
| Bacterial_alpha-L-rhamnosidase_6_hairpin_glycosidase_domain |  |  | 1 |  |  | 1 |  |  | 1 |
| Bacterial_alpha-L-rhamnosidase_concanavalin-like_domain |  |  | 1 |  |  | 1 |  |  | 1 |
| Bacterial_protein_of_unknown_function_(DUF882) |  |  | 1 |  |  | 1 |  |  | 1 |
| Bacterial_toxin_homologue_of_phage_lysozyme,_C-term |  |  | 1 |  |  |  |  |  |  |
| Chitin_binding_Peritrophin-A_domain |  |  | 1 |  |  | 1 |  |  |  |
| Chitin_recognition_protein |  |  | 1 |  |  | 1 |  |  | 1 |
| Chitinase_class_I | 24 | 1 | 1 | 24 | 1 | 1 | 24 | 1 | 1 |
| Cutinase |  |  | 1 |  |  |  |  |  |  |
| DDE_superfamily_endonuclease |  |  | 1 |  |  | 1 |  |  | 1 |
| Lysozyme_like_domain |  | 1 | 1 |  |  |  |  |  |  |
| ribonuclease |  |  | 1 |  |  | 1 |  |  |  |
| Ribosomal_protein_L7Ae/L30e/S12e/Gadd45_family |  |  | 1 |  |  | 1 |  |  |  |
| Serine_carboxypeptidase |  |  | 1 |  |  | 1 |  |  |  |
| Transposase |  |  | 1 |  |  | 1 |  |  | 1 |
| von_Willebrand_factor_type_A_domain |  |  | 1 |  |  | 1 |  |  |  |
| Domain_of_unknown_function_(DUF7029) | 1 |  |  |  |  |  |  |  |  |
| Domain_of_unknown_function_(DUF7223) | 1 |  |  |  |  |  |  |  |  |
| DZIP3/_hRUL138-like_HEPN | 1 |  |  | 1 |  |  | 1 |  |  |
| Fungal_cellulose_binding_domain | 2 |  |  |  |  |  |  |  |  |
| Lipase |  |  |  |  |  |  |  | 1 |  |
| Mannosyl-glycoprotein_endo-beta-N-acetylglucosaminidase |  |  |  |  | 1 |  |  | 1 |  |

### Cysteine-rich repeats often hallmark cross-domain horizontal gene transfer in eukaryotes

Our analyses suggest that CysRReps may serve as recognizable markers of horizontally acquired protein parts in eukaryotic genomes. In the GL protein of *N. californiae*, enzymatic domains display strong similarity to bacterial and bacteriophage sequences, whereas the intervening CysRReps lack detectable homologs in prokaryotes. This pattern suggests that foreign coding regions integrated into a pre-existing eukaryotic sequence context and acquired CysRReps during assimilation.

We propose two alternative HGT scenarios associated with CysRReps. The first involves a transfer of a bacterial chromosomal fragment to a bacteriophage via transduction, followed by nonspecific integration of bacteriophage DNA into the eukaryotic chromosome (Fig. 7A). The second scenario entails direct transfer of bacterial DNA into the eukaryotic genome (Fig. 7B), as has been proposed for other eukaryotic lineages (Hespeels et al. 2014; Lacroix and Citovsky 2016; Bininda-Emonds et al. 2016). The genomic context of the closest bacterial homologs further supports a phage-mediated transfer scenario, as the homologous genes reside within prophage regions and occur near recombination-associated elements such as integrases and recombination sites (Thorpe and Smith 1998; Julien 2003). These observations support a model in which bacteriophages act as vectors that facilitate the movement of bacterial DNA into eukaryotic genomes.

**Fig. 7.**
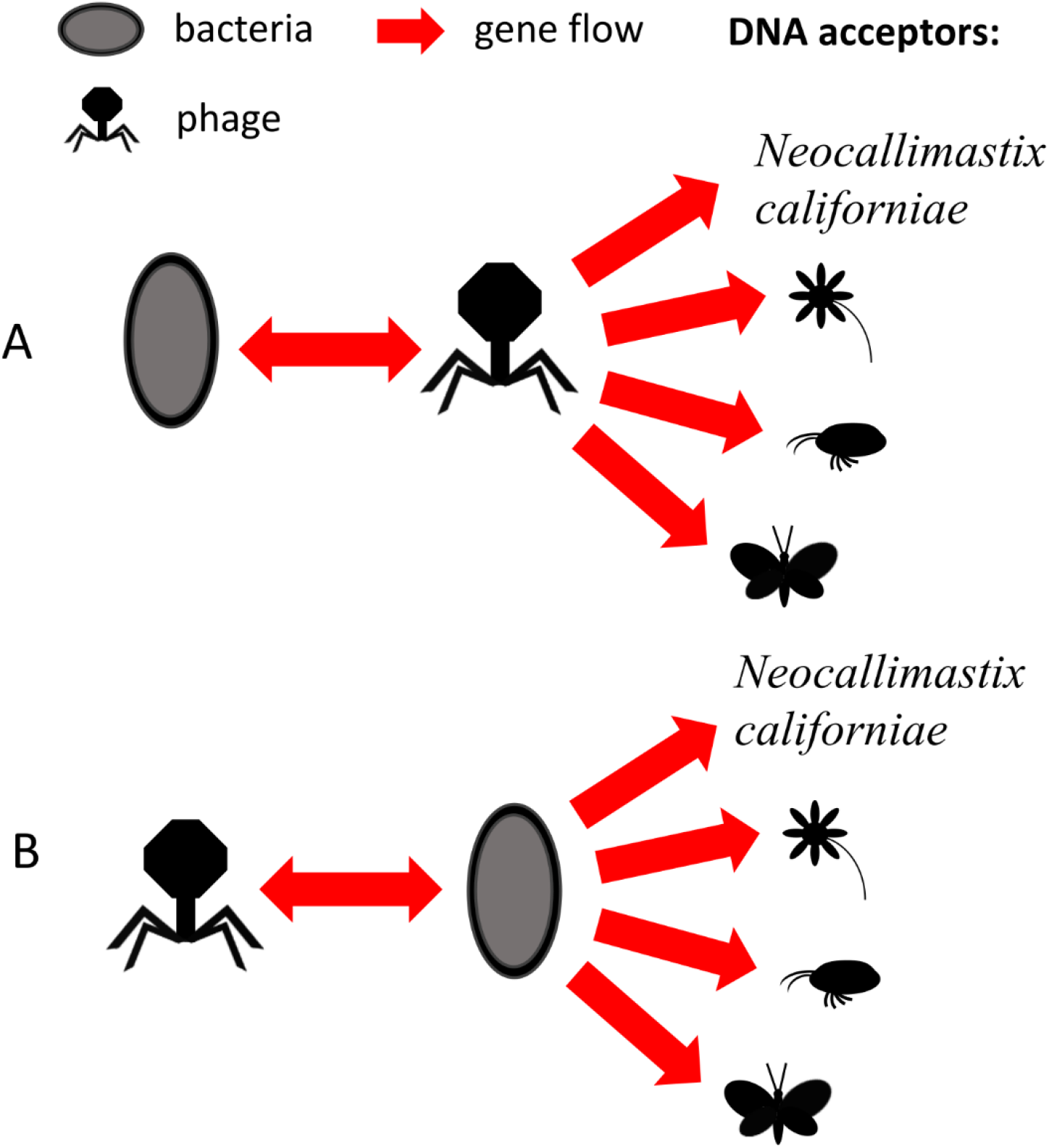
Two scenarios of gene transfer between prokaryotes and eukaryotes. **A** DNA exchange between phages and bacteria, and direct DNA transfer from bacteria to eukaryotes **B** phage as a mediator of DNA transfer between bacteria and eukaryotes

Beyond the specific case of *N. californiae*, the broader dataset analyzed here indicates that proteins containing CysRReps occur across diverse eukaryotic lineages and frequently include sequences with strong similarity to bacterial, archaeal, or viral proteins. The taxonomic distribution of eukaryotic proteins is consistent with HGT frequencies previously reported in fungi, rotifers, and other eukaryotes (Eyres et al. 2015; Liu et al. 2024). The dominance of Actinomycetes among potential bacterial partners also agrees with earlier studies showing that this group contributes a substantial fraction of bacterial donors in fungal HGT events (Liu et al. 2024). Viral homologs detected in our dataset are overwhelmingly restricted to Caudoviricetes, suggesting that this group of bacteriophages may frequently mediate the transfer of bacterial genes across domain boundaries. Increasing evidence indicates that bacteriophages can interact with eukaryotic cells more extensively than previously recognized, including the ability to enter eukaryotic cells or deliver DNA into the nucleus (Redrejo-Rodríguez et al. 2012; Lehti et al. 2017).

The functional role of CysRReps remains unclear. In the GL protein, as well as in other examples identified here, these repeats appear to precede horizontally acquired enzymatic domains. They therefore may represent eukaryotic modules that became associated with foreign coding regions after integration. Such architecture could facilitate the assimilation of transferred genes by providing regulatory or structural context, for example, by enabling promoter capture or the formation of new multidomain proteins. Similar processes have been observed during experimental gene transfer events and in natural cross-domain HGT (Husnik and McCutcheon 2018; Li and Bock 2019; Callens et al. 2021). An alternative possibility is that CysRReps mark genomic regions that were previously involved in bacteriophage integration events, as proposed earlier for other eukaryotic genomes (Daugavet et al. 2019). In either case, the consistent association between CysRReps and prokaryote-derived protein parts suggests that these motifs may represent useful indicators of past horizontal gene transfer events and the subsequent evolutionary assimilation of foreign genes into eukaryotic genomes.

## Conclusions

We identify a consistent association between CysRReps in eukaryotic proteins and adjacent sequences of likely prokaryotic origin. Although the mechanistic role of these repeats remains unclear, their recurrent proximity to bacterial-, archaeal-, and viral-like domains suggests that they mark genomic regions where foreign DNA has integrated and become assimilated into eukaryotic proteins. Analysis of a large and taxonomically diverse dataset shows that proteins with this architecture occur across multiple eukaryotic lineages, indicating that such events are widespread rather than restricted to specific taxa. The strong representation of bacteriophage-related sequences among the closest homologs further suggests that phages may act as intermediaries linking prokaryotic gene pools with eukaryotic genomes. Together, these results highlight CysRReps as potential indicators of past cross-domain horizontal gene transfer and provide a new framework for identifying and investigating gene exchange across the tree of life.

## Abbreviations

HGT: Horizontal Gene Transfer
GL: Glucosamidase-Like
GH: Glycosyl Hydrolase
CysRRep: Cysteine-Rich Repeat

## Statements and Declarations

### Ethics approval and consent to participate

Not applicable

### Consent for publication

Not applicable

### Availability of data and materials

All data generated or analyzed during this study are included in this published article and its supplementary information files.

## Authors’ contributions

MD conceived and designed the study, performed the analyses, and drafted the manuscript. SM contributed to coding and data processing. VD curated the AttP collection, NE and MR-B provided critical feedback on the study, and contributed to funding. MD wrote the manuscript with contributions from all the co-authors.

## Acknowledgements

The authors thank Professor Olga Podgornaya for her years of support and inspiration.

## Supplementary Information

**Fig. S1.**
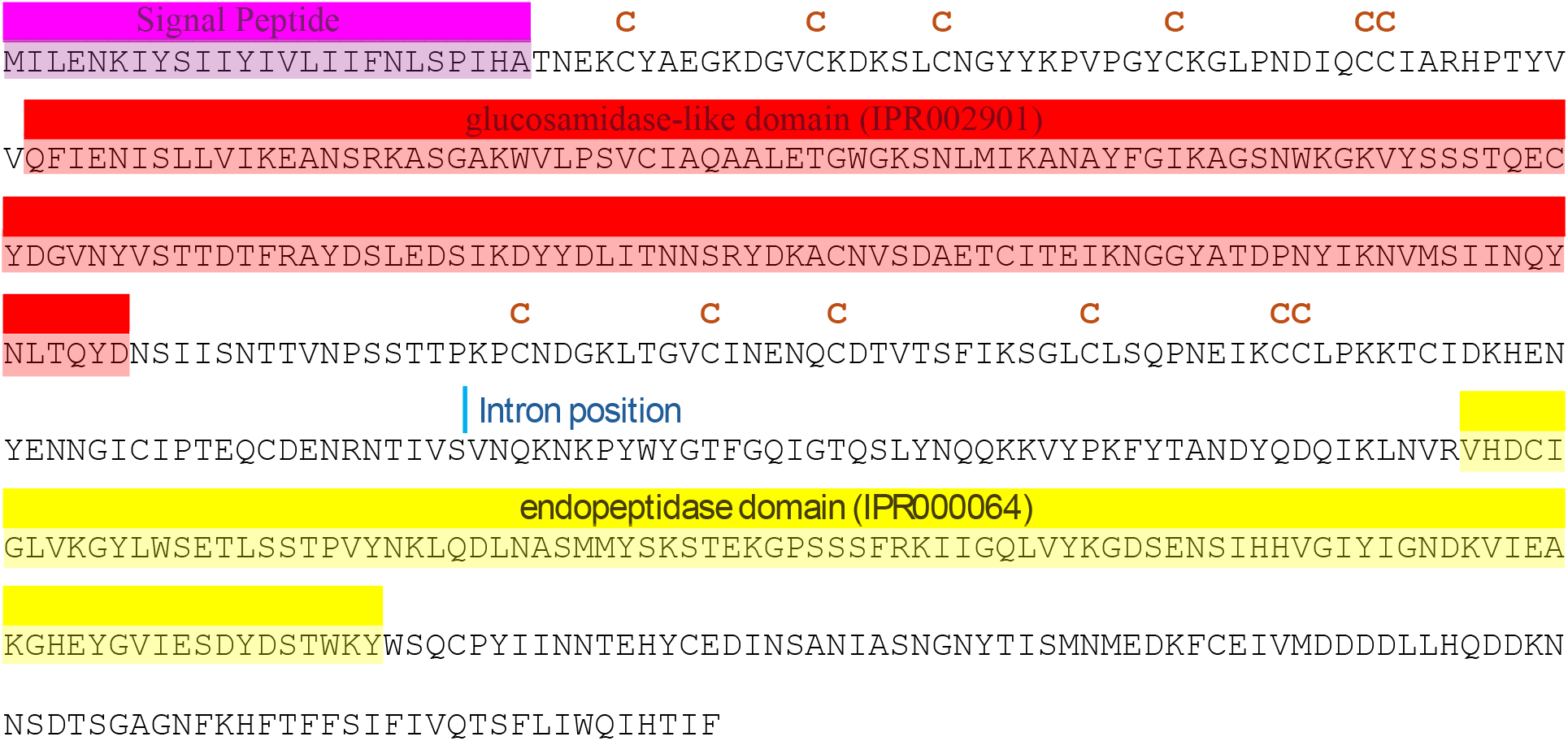
Features and primary sequence of glucosaminidase-domain-containing protein of *Neocallimastix californiae* (ID: ORY21964.1). Brown “C” indicate positions of conservative cysteines in cysteine-rich repeats.

**Fig. S2.**
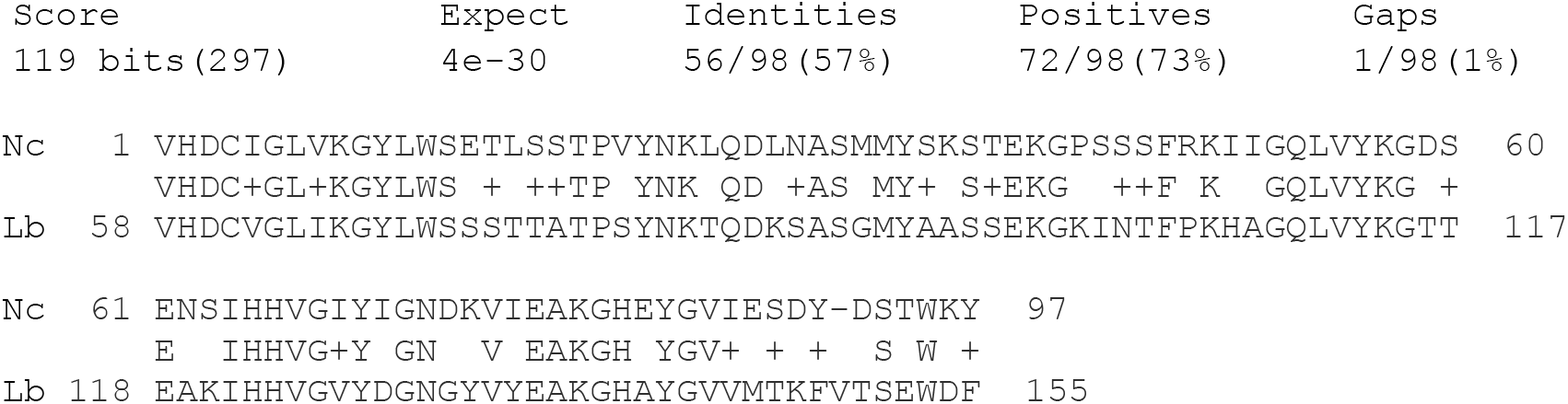
Alignment of (Nc) *N. californiae* sequence corresponding to predicted endopeptidase domain with (Lb) *Lachnospiraceae bacterium* NlpC/P60 family protein (ID: MCD8119760.1)

**Fig. S3.**
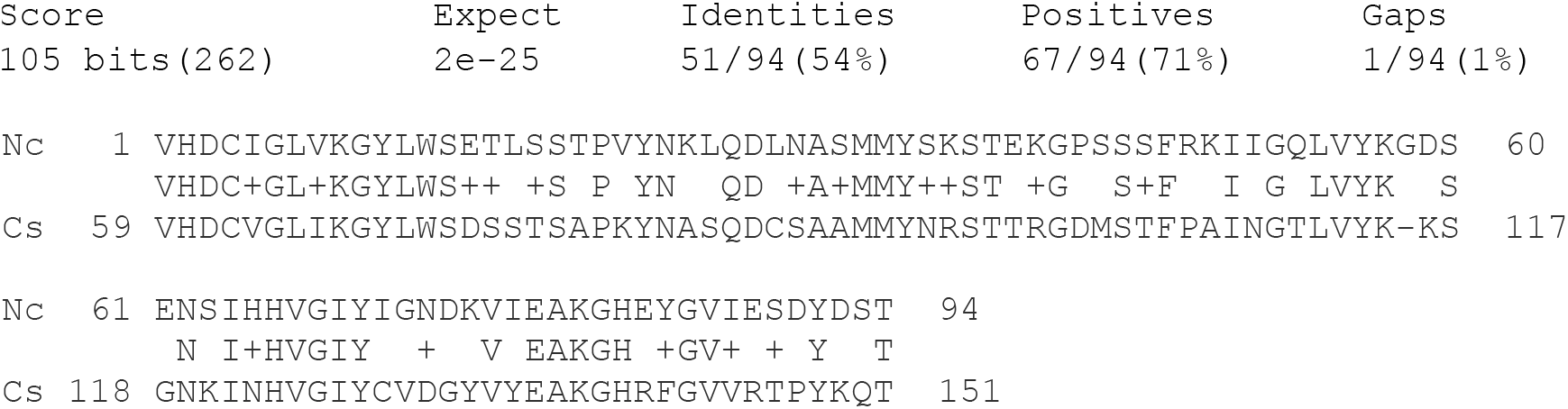
Alignment of *N. californiae* (Nc) sequence corresponding to predicted endopeptidase domain with CW7 repeat protein of Caudoviricetes sp. (Cs) protein (ID: DAT38862.1)

**Supplementary File 1**

Database of bacteriophage recombination sites extracted from GenBank records (.fasta).

**Supplementary File 2**

Alignment of five cysteine-rich repeats of *N. californiae* proteins (.fasta)

**Supplementary File 3**

Script for JackHMMER and mmseqs2 searches (.bash).

**Supplementary File 4**

Script for extraction of protein regions (.R).

**Supplementary File 5**

Script for mmseqs2 searche of protein regions against the Pfam database (.bash).

**Supplementary File 6**

Prophage region of Lachnospiraceae bacterium isolate C.4_2 harboring the closest bacterial homolog of GL protein endopeptidase domain (.txt).

**Supplementary File 7**

Results of the JackHMMER search of cysteine-rich repeat alignment against the UniProtKB database (.txt).

**Supplementary File 8**

N-terminal regions adjacent to cysteine-rich repeats in eukaryotic proteins (.fasta).

**Supplementary File 9**

C-terminal regions adjacent to cysteine-rich repeats in eukaryotic proteins (.fasta).

**Supplementary File 10**

Regions located between cysteine-rich repeats in eukaryotic proteins (.fasta).

**Supplementary Table T1**

BLASTp results of GL protein search against eukaryotic proteins (.xlsx).

**Supplementary Table T2**

BLASTp results of GL protein search against bacterial proteins (.xlsx).

**Supplementary Table T3**

BLASTp results of GL protein search against viral proteins (.xlsx).

**Supplementary Table T4**

Bacterial hits extracted from the JackHMMER search of the cysteine-rich repeat alignment against UniProtKB (.xlsx).

**Supplementary Table T5**

Results of mmseqs2 searches of eukaryotic proteins regions against archaeal, bacterial and viral sequences (.xlsx).

