## Supplementary figure S1 for "Cysteine-rich repeats trace past horizontal gene transfers in eukaryotes"

Signal Peptide
C C C C CC
MILENKIYSIIYIVLIIFNLSPIHATNEKCYAEGKDGVCKDKSLCNGYYKPVPGYCKGLPNDIQCCIARHPTYV

glucosamidase-like domain (IPR002901)
VQFIENISLLVIKEANSRKASGAKWVLPSVCIAQAALETGWGKSNLMIKANAYFGIKAGSNWKGKVYSSSTQEC

YDGVNYVSTTDTFRAYDSLEDSIKDYYDLITNNSRYDKACNVSDAETCITEIKNGGYATDPNYIKNVMSIINQY

C C C C CC
NLTQYDNSIISNTTVNPSSTTPKPCNDGKLTGVCINENQCDTVTSFIKSGLCLSQPNEIKCCLPKKTCIDKHEN

Intron position
YENNGICIPTEQCDENRNTIVSVNQKNKPYWYGTFGQIGTQSLYNQQKKVYPKFYTANDYQDQIKLNVRVHDCI

endopeptidase domain (IPR000064)
GLVKGYLWSETLSSTPVYNKLQDLNASMMYSKSTEKGPSSSFRKIIGQLVYKGDSENSIHHVGIYIGNDKVIEA

KGHEYGVIESDYDSTWKYWSQCPYIINNTEHYCEDINSANIASNGNYTISMNMEDKFCEIVMDDDDLLHQDDKN

NSDTSGAGNFKHFTFFSIFIVQTSFLIWQIHTIF

**Fig S1**

Features and primary sequence of glucosaminidase-domain-containing protein of *Neocallimastix californiae* (ID: ORY21964.1). Brown “C” indicate positions of conservative cysteines in cysteine-rich repeats.
