## Supplementary figure S2 for "Cysteine-rich repeats trace past horizontal gene transfers in eukaryotes"

| Score | Expect | Identities | Positives | Gaps |
|---|---|---|---|---|
| 119 bits(297) | 4e-30 | 56/98(57%) | 72/98(73%) | 1/98(1%) |

```
Nc   1 VHDCIGLVKGYLWSETLSSTPVYNKLQDLNASMMYSKSTEKGPSSSFRKIIGQLVYKGDS  60
       VHDC+GL+KGYLWS + ++TP YNK QD +AS MY+ S+EKG  ++F K  GQLVYKG +
Lb  58 VHDCVGLIKGYLWSSSTTATPSYNKTQDKSASGMYAASSEKGKINTFPKHAGQLVYKGTT  117

Nc  61 ENSIHHVGIYIGNDKVIEAKGHEYGVIESDY-DSTWKY  97
       E  IHHVG+Y GN  V EAKGH YGV+ + +  S W +
Lb 118 EAKIHHVGVYDGNGYVYEAKGHAYGVVMTKFVTSEWDF  155
```

**Fig S2**

Alignment of (Nc) *N. californiae* sequence corresponding to predicted endopeptidase domain with (Lb) *Lachnospiraceae bacterium* NlpC/P60 family protein (ID: MCD8119760.1)
