## Supplementary figure S3 for "Cysteine-rich repeats trace past horizontal gene transfers in eukaryotes"

```
Score               Expect     Identities      Positives       Gaps
105 bits(262)       2e-25      51/94(54%)      67/94(71%)      1/94(1%)

Nc   1 VHDCIGLVKGYLWSETLSSTPVYNKLQDLNASMMYSKSTEKGPSSSFRKIIGQLVYKGDS  60
       VHDC+GL+KGYLWS++ +S P YN  QD +A+MMY++ST +G  S+F  I G LVYK  S
Cs  59 VHDCVGLIKGYLWSDSSTSAPKYNASQDCSAAMMYNRSTTRGDMSTFPAINGTLVYK-KS  117

Nc  61 ENSIHHVGIYIGNDKVIEAKGHEYGVIESDYDST  94
        N I+HVGIY   +  V EAKGH +GV+ + Y  T
Cs 118 GNKINHVGIYCVDGYVYEAKGHRFGVVRTPYKQT  151
```

**Fig S3**

Alignment of *N. californiae* (Nc) sequence corresponding to predicted endopeptidase domain with CW7 repeat protein of Caudoviricetes sp. (Cs) protein (ID: DAT38862.1)
