## Supplementary file 2 for "Cysteine-rich repeats trace past horizontal gene transfers in eukaryotes"

; Journal of Molecular Evolution													
; Cysteine-rich repeats trace past horizontal gene transfers in eukaryotes												
; Maria Daugavet1,2, Viktoria Dikaya3, Natella Enukashvily3, Stas Malavin1,4, Maxim Rubin-Blum1,2.
; 1.Biology Department, National Institute of Oceanography, Israel Oceanographic and Limnological Research (IOLR), Haifa 3108000, Israel	
; 2.Department of Marine Biology, Leon H. Charney School of Marine Sciences, University of Haifa, Haifa 3498838, Israel	
; 3.Laboratory of the Non-Coding DNA Study, Institute of Cytology, Russian Academy of Sciences, St. Petersburg 194064, Russia	
; 4.Department of Environmental Hydrology and Microbiology, Zuckerberg Institute for Water Research, The Jacob Blaustein Institutes for Desert Research, Ben-Gurion University of the Negev, Sde Boker 8499000, Israel	

>ORY85279.1_1
LNEKCTNYHGLSGECKDISQSCDTGINFIIPYLCEGKDN-IKCC
>ORY85279.1_2
PKNTCNDDRGNQGFCIP-SSECNTNYHNIVSEKCASGDNTIKCC
>ORY85279.1_3
NSSPCSD-GNLAGKCIN-ENLCDKNTHFIKAGLCLNQPNEIKCC
>ORY21964.1_1
TNEKCYA-EGKDGVCKD-KSLCN-GYYKPVPGYCKGLPNDIQCC
>ORY21964.1_2
TPKPCND-GKLTGVCIN-ENQCDTVTSFIKSGLCLSQPNEIKCC
