## Supplementary file 4 for "Cysteine-rich repeats trace past horizontal gene transfers in eukaryotes"

# Journal of Molecular Evolution
# Cysteine-rich repeats trace past horizontal gene transfers in eukaryotes
# Maria Daugavet1,2, *, Viktoria Dikaya3, Natella Enukashvily3, Stas Malavin1,4, Maxim Rubin-Blum1,2
# 1. Biology Department, National Institute of Oceanography, Israel Oceanographic and Limnological Research (IOLR), Haifa 3108000, Israel
# 2. Department of Marine Biology, Leon H. Charney School of Marine Sciences, University of Haifa, Haifa 3498838, Israel
# 3. Laboratory of the Non-Coding DNA Study, Institute of Cytology, Russian Academy of Sciences, St. Petersburg 194064, Russia
# 4. Department of Environmental Hydrology and Microbiology, Zuckerberg Institute for Water Research, The Jacob Blaustein Institutes for Desert Research, Ben-Gurion University of the Negev, Sde Boker 8499000, Israel

#Script for extraction of protein regions

library(dplyr)

tmp <- read.csv('tmp.csv', header = F)[ ,1:22]

# Write a file with repeat coordinates for the final output
tmp %>% mutate(Rep = paste(V20, V21, sep = ":")) %>%
	group_by(V4, V1) %>% summarize(Rep = paste(Rep, collapse = ",")) %>%
	write.table(file = 'res_rep', row.name = F, col.names = F, quote = F, sep = '\t')

sliced <- data.frame(target = NA, start = NA, end = NA, segment = NA)

for(id in unique(tmp$V1)) {
	message(id, ' (', match(id, unique(tmp$V1)), '/', length(unique(tmp$V1)), ')')
	tmpp <- tmp[tmp$V1 %in% id, ]
	if (nrow(tmpp) > 1) {
		for (i in 2:nrow(tmpp)) {
			sliced[nrow(sliced)+1,] <- c(tmpp[i,1], tmpp[i-1,21], tmpp[i,20], tmpp[i,10])
		}
	}
	sliced[nrow(sliced)+1,] <- c(tmpp[1,1], 1, min(tmpp[,20]), 'b')
	sliced[nrow(sliced)+1,] <- c(tmpp[nrow(tmpp),1], max(tmpp[,21]),
								 tmpp[nrow(tmpp),3], 'e')
}

sliced <- sliced[!is.na(sliced$segment) & sliced$segment != 1, ]

sliced$start <- as.numeric(sliced$start)
sliced$end <- as.numeric(sliced$end)

sliced %<>% group_by(target, segment) %>% summarize(start = min(start), end = min(end)) %>%
	mutate(len = end - start) %>%
	filter(len >= 100)

sliced %>% filter(segment == 'e') %>%
	write.table(file = 'sliced_e.tsv', quote = F, row.names = F)

sliced %>% filter(segment == 'b') %>%
	write.table(file = 'sliced_b.tsv', quote = F, row.names = F)

sliced %>% filter(segment != 'b' & segment != 'e') %>%
	write.table(file = 'sliced_i.tsv', quote = F, row.names = F)
