## Supplementary file 6 for "Cysteine-rich repeats trace past horizontal gene transfers in eukaryotes"

**Journal of Molecular Evolution**

**Phaster (**[**https://phastest.ca/**](https://phastest.ca/)**) output of prophage region of Lachnospiraceae bacterium isolate C.4_2**

**gi|00000000|ref|JAJQBQ010000044.1| MAG: Lachnospiraceae bacterium isolate C.4_2 174419,**

**gc%: 47.61%**

Hits against Virus and Prophage Database - purple

Hits against Bacterial Database or GenBank File – pink

Hit to *N. californiae* at position 98280..99080 – bold

Total 96 CDS

| **#** | **CDS Position** | **BLAST Hit** | **E-Value** |
| --- | --- | --- | --- |
| 1 | 52845..52869 | attL | 0.0 |
| 2 | complement(53022..54239) | PHAGE_Faecal_FP_oengus_NC_047916: methyltransferase type 11; LUE29_09450; phage(gi100001) | 3.61e-131 |
| 3 | complement(54246..54449) | hypothetical protein; LUE29_09455 | 0.0 |
| 4 | 54508..54672 | hypothetical protein; LUE29_09460 | 0.0 |
| 5 | 54795..55106 | DUF6431 domain-containing protein; LUE29_09465 | 0.0 |
| 6 | 55352..55425 | tRNA | 0.0 |
| 7 | 55723..55971 | PHAGE_Faecal_FP_oengus_NC_047916: hypothetical protein; LUE29_09475; phage(gi100002) | 4.73e-35 |
| 8 | 55971..56393 | PHAGE_Faecal_FP_oengus_NC_047916: hypothetical protein; LUE29_09480; phage(gi100003) | 1.76e-24 |
| 9 | 56399..56575 | hypothetical protein; LUE29_09485 | 0.0 |
| 10 | 56645..57019 | hypothetical protein; LUE29_09490 | 0.0 |
| 11 | 57169..57387 | hypothetical protein; LUE29_09495 | 0.0 |
| 12 | 57436..57807 | hypothetical protein; LUE29_09500 | 0.0 |
| 13 | 57959..58255 | DNA-directed RNA polymerase I; LUE29_09505 | 0.0 |
| 14 | 58323..58682 | PHAGE_Faecal_FP_oengus_NC_047916: hypothetical protein; LUE29_09510; phage(gi100009) | 3.13e-44 |
| 15 | 58737..58916 | hypothetical protein; LUE29_09515 | 0.0 |
| 16 | 59016..59225 | CTP synthase; LUE29_09520 | 0.0 |
| 17 | 59299..59370 | tRNA | 0.0 |
| 18 | 59413..60168 | PHAGE_Faecal_FP_oengus_NC_047916: hypothetical protein; LUE29_09530; phage(gi100015) | 1.08e-30 |
| 19 | 60223..60402 | hypothetical protein; LUE29_09535 | 0.0 |
| 20 | 60450..61223 | PHAGE_Gordon_Untouchable_NC_048823: putative single-stranded DNA binding protein; LUE29_09540; phage(gi100050) | 7.89e-61 |
| 21 | 61293..61556 | hypothetical protein; LUE29_09545 | 0.0 |
| 22 | 61645..61842 | hypothetical protein; LUE29_09550 | 0.0 |
| 23 | 61842..62471 | PHAGE_Gordon_Untouchable_NC_048823: hypothetical protein; LUE29_09555; phage(gi100059) | 8.62e-30 |
| 24 | 62481..62726 | PHAGE_Klebsi_vB_KpnM_KpV79_NC_042041: hypothetical protein; LUE29_09560; phage(gi100013) | 4.66e-17 |
| 25 | 62751..63701 | PHAGE_Faecal_FP_oengus_NC_047916: hypothetical protein; LUE29_09565; phage(gi100022) | 8.77e-59 |
| 26 | 63884..64036 | PHAGE_Faecal_FP_oengus_NC_047916: hypothetical protein; LUE29_09570; phage(gi100024) | 3.88e-15 |
| 27 | 64037..64588 | PHAGE_Faecal_FP_oengus_NC_047916: hypothetical protein; LUE29_09575; phage(gi100025) | 3.26e-16 |
| 28 | 64588..65175 | PHAGE_Faecal_FP_oengus_NC_047916: hypothetical protein; LUE29_09580; phage(gi100026) | 1.47e-44 |
| 29 | 65189..65569 | hypothetical protein; LUE29_09585 | 0.0 |
| 30 | 65571..69608 | PHAGE_Faecal_FP_oengus_NC_047916: hypothetical protein; LUE29_09590; phage(gi100028) | 0.0 |
| 31 | 69629..69811 | hypothetical protein; LUE29_09595 | 0.0 |
| 32 | 69813..70283 | PHAGE_Gordon_Avazak_NC_048822: portal protein; LUE29_09600; phage(gi100069) | 9.65e-47 |
| 33 | 70309..70596 | hypothetical protein; LUE29_09605 | 0.0 |
| 34 | 70626..70988 | hypothetical protein; LUE29_09610 | 0.0 |
| 35 | 71005..71217 | hypothetical protein; LUE29_09615 | 0.0 |
| 36 | 71266..71481 | hypothetical protein; LUE29_09620 | 0.0 |
| 37 | 71478..72692 | PHAGE_Faecal_FP_oengus_NC_047916: hypothetical protein; LUE29_09625; phage(gi100035) | 0.0 |
| 38 | 72686..72880 | hypothetical protein; LUE29_09630 | 0.0 |
| 39 | 72900..73121 | hypothetical protein; LUE29_09635 | 0.0 |
| 40 | 73123..73386 | hypothetical protein; LUE29_09640 | 0.0 |
| 41 | 73389..73613 | hypothetical protein; LUE29_09645 | 0.0 |
| 42 | 74147..74455 | PHAGE_Butyri_Arawn_NC_048848: hypothetical protein; LUE29_09650; phage(gi100031) | 6.42e-07 |
| 43 | 74446..74658 | hypothetical protein; LUE29_09655 | 0.0 |
| 44 | 74669..75001 | PHAGE_Faecal_FP_Epona_NC_047910: hypothetical protein; LUE29_09660; phage(gi100017) | 2.42e-27 |
| 45 | 74998..75750 | PHAGE_Clostr_phiCTP1_NC_014457: hypothetical protein; LUE29_09665; phage(gi304360730) | 4.27e-23 |
| 46 | 75731..75934 | hypothetical protein; LUE29_09670 | 0.0 |
| 47 | 75956..76279 | PHAGE_Clostr_phiCTC2B_NC_030951: hypothetical protein; LUE29_09675; phage(gi100016) | 6.68e-05 |
| 48 | 76287..76454 | hypothetical protein; LUE29_09680 | 0.0 |
| 49 | 76473..76754 | hypothetical protein; LUE29_09685 | 0.0 |
| 50 | 76687..77055 | PHAGE_Eggert_PMBT5_NC_048022: hypothetical protein; LUE29_09690; phage(gi100014) | 7.08e-08 |
| 51 | 77048..77203 | hypothetical protein; LUE29_09695 | 0.0 |
| 52 | 77233..77427 | hypothetical protein; LUE29_09700 | 0.0 |
| 53 | 77433..77612 | hypothetical protein; LUE29_09705 | 0.0 |
| 54 | 78329..78826 | DUF1232 domain-containing protein; LUE29_09710 | 0.0 |
| 55 | 79084..79374 | PHAGE_Faecal_FP_oengus_NC_047916: tail assembly protein; LUE29_09715; phage(gi100053) | 8.08e-36 |
| 56 | 79346..80023 | PHAGE_Gordon_OhMyWard_NC_048816: hypothetical protein; LUE29_09720; phage(gi100084) | 3.01e-56 |
| 57 | 80028..82994 | PHAGE_Faecal_FP_oengus_NC_047916: minor tail protein; LUE29_09725; phage(gi100055) | 0.0 |
| 58 | 83034..83276 | PHAGE_Faecal_FP_oengus_NC_047916: minor tail protein; LUE29_09730; phage(gi100056) | 2.38e-23 |
| 59 | 83605..83943 | PHAGE_Gordon_Untouchable_NC_048823: hypothetical protein; LUE29_09735; phage(gi100010) | 6.55e-38 |
| 60 | 83933..84349 | PHAGE_Gordon_Chikenjars_NC_048810: hypothetical protein; LUE29_09740; phage(gi100012) | 5.87e-52 |
| 61 | 84367..84714 | PHAGE_Faecal_FP_oengus_NC_047916: hypothetical protein; LUE29_09745; phage(gi100062) | 2.95e-39 |
| 62 | 84718..85251 | PHAGE_Faecal_FP_oengus_NC_047916: portal protein; LUE29_09750; phage(gi100069) | 6.38e-15 |
| 63 | 85255..85758 | PHAGE_Faecal_FP_oengus_NC_047916: portal protein; LUE29_09755; phage(gi100069) | 1.93e-10 |
| 64 | 85780..87525 | PHAGE_Gordon_Secretariat_NC_048876: hypothetical protein; LUE29_09760; phage(gi100013) | 0.0 |
| 65 | 87816..88940 | PHAGE_Rhodoc_ReqiPepy6_NC_023735: gp018; LUE29_09765; phage(gi593779805) | 9.10e-08 |
| 66 | 89003..89161 | hypothetical protein; LUE29_09770 | 0.0 |
| 67 | 89238..89714 | PHAGE_Faecal_FP_oengus_NC_047916: portal protein; LUE29_09775; phage(gi100069) | 2.38e-10 |
| 68 | 89737..90285 | PHAGE_Faecal_FP_oengus_NC_047916: portal protein; LUE29_09780; phage(gi100069) | 2.97e-10 |
| 69 | 90296..91396 | PHAGE_Gordon_Kenosha_NC_048793: hypothetical protein; LUE29_09785; phage(gi100016) | 9.07e-10 |
| 70 | 91393..91581 | hypothetical protein; LUE29_09790 | 0.0 |
| 71 | 91601..92863 | PHAGE_Faecal_FP_oengus_NC_047916: terminase large subunit; LUE29_09795; phage(gi100070) | 0.0 |
| 72 | 92853..94553 | PHAGE_Gordon_Duffington_NC_048750: hypothetical protein; LUE29_09800; phage(gi100022) | 0.0 |
| 73 | 94583..94906 | PHAGE_Arthro_Gordon_NC_041952: hypothetical protein; LUE29_09805; phage(gi100014) | 5.69e-35 |
| 74 | 94977..95195 | PHAGE_Gordon_Secretariat_NC_048876: hypothetical protein; LUE29_09810; phage(gi100022) | 1.92e-19 |
| 75 | 95215..95865 | PHAGE_Strept_Rima_NC_041889: hypothetical protein; LUE29_09815; phage(gi100017) | 2.99e-88 |
| 76 | 95891..96628 | PHAGE_Gordon_Avazak_NC_048822: hypothetical protein; LUE29_09820; phage(gi100022) | 1.39e-67 |
| 77 | 96694..97098 | PHAGE_Faecal_FP_oengus_NC_047916: hypothetical protein; LUE29_09825; phage(gi100076) | 1.42e-37 |
| 78 | 97126..97671 | PHAGE_Faecal_FP_oengus_NC_047916: hypothetical protein; LUE29_09830; phage(gi100077) | 7.57e-60 |
| 79 | 97674..98039 | hypothetical protein; LUE29_09835 | 0.0 |
| 80 | 98036..98278 | hypothetical protein; LUE29_09840 | 0.0 |
| **81** | **98280..99080** | **NlpC/P60 family protein; LUE29_09845** | **0.0** |
| 82 | 99091..99468 | PHAGE_Faecal_FP_oengus_NC_047916: single-stranded-DNA-specific exonuclease; LUE29_09850; phage(gi100078) | 1.54e-39 |
| 83 | 99465..104705 | PHAGE_Arthro_Circum_NC_041948: hypothetical protein; LUE29_09855; phage(gi100022) | 4.51e-166 |
| 84 | 104716..105562 | hypothetical protein; LUE29_09860 | 0.0 |
| 85 | 105573..107183 | PHAGE_Arthro_Circum_NC_041948: hypothetical protein; LUE29_09865; phage(gi100022) | 1.51e-11 |
| 86 | 107184..107987 | PHAGE_Faecal_FP_oengus_NC_047916: hypothetical protein; LUE29_09870; phage(gi100081) | 8.47e-29 |
| 87 | 107991..111536 | PHAGE_Faecal_FP_oengus_NC_047916: putative replicative DNA helicase; LUE29_09875; phage(gi100080) | 4.38e-24 |
| 88 | 111562..111939 | hypothetical protein; LUE29_09880 | 0.0 |
| 89 | 111968..112807 | PHAGE_Lactob_JCL1032_NC_019456: structural protein; LUE29_09885; phage(gi418489132) | 3.82e-63 |
| 90 | 112840..113670 | PHAGE_Lactob_JCL1032_NC_019456: structural protein; LUE29_09890; phage(gi418489132) | 1.00e-67 |
| 91 | 113727..114035 | PHAGE_Lactob_JCL1032_NC_019456: hypothetical protein; LUE29_09895; phage(gi418489133) | 6.50e-16 |
| 92 | 114051..114194 | PHAGE_Butyri_Arawn_NC_048848: hypothetical protein; LUE29_09900; phage(gi100020) | 4.63e-05 |
| 93 | 114191..114679 | hypothetical protein; LUE29_09905 | 0.0 |
| 94 | 114891..116180 | PHAGE_Faecal_FP_Taranis_NC_047914: terminase small subunit; LUE29_09910; phage(gi100071) | 1.42e-31 |
| 95 | 116149..116454 | PHAGE_Faecal_FP_Brigit_NC_047909: hypothetical protein; LUE29_09915; phage(gi100064) | 4.59e-19 |
| 96 | 116603..116627 | attR | 0.0 |
